# Adeno-associated virus mediated gene delivery: Implications for scalable *in vitro* and *in vivo* cardiac optogenetic models

**DOI:** 10.1101/183319

**Authors:** Christina M. Ambrosi, Gouri Sadananda, Aleksandra Klimas, Emilia Entcheva

## Abstract

**Aims:** Adeno-associated viruses (AAVs) provide advantages in long-term, cardiac-specific gene expression. However, AAV serotype specificity data is lacking in cardiac models relevant to optogenetics. We aimed to identify the optimal AAV serotype (1, 6, or 9) in pursuit of scalable rodent and human models for cardiac optogenetics and elucidate the mechanism of virus uptake.

**Methods:** *In vitro* syncytia of primary neonatal rat ventricular cardiomyocytes (NRVMs) and human induced pluripotent stem cell-derived cardiomyocytes (hiPSC-CMs) were infected with AAVs 1, 6, and 9 containing the transgene for eGFP or channelrhodopsin-2 (ChR2) fused to mCherry. *In vivo* adult rats were intravenously injected with AAV1 and 9 containing ChR2-mCherry.

**Results:** Transgene expression profiles of rat and human cells *in vitro* revealed that AAV1 and 6 significantly outperformed AAV9. In contrast, systemic delivery of AAV9 in adult rat hearts yielded significantly higher levels of ChR2-mCherry expression and optogenetic responsiveness. We tracked the mechanism of virus uptake to purported receptor-mediators for AAV 1/6 (cell surface sialic acid) and AAV9(37/67kDa laminin receptor, LamR). *In vitro* desialylation of NRVMs and hiPSC-CMs with neuraminidase significantly decreased AAV1,6-mediated gene expression, but interestingly, desialylation of hiPSC-CMs increased AAV9-mediated expression. In fact, only very high viral doses of AAV9-ChR2-mCherry, combined with neuraminidase treatment yielded consistent optogenetic responsiveness in hiPSC-CMs. Differences between the *in vitro* and *in vivo* performance of AAV9 could be correlated to robust LamR expression in the adult and neonatal rat hearts, but no expression *in vitro* in cultured cells. The dynamic nature of LamR expression and its dependence on environmental factors was further corroborated in intact adult human ventricular tissue slices.

**Conclusion:** The combined transgene expression and cell surface receptor data may explain the preferential efficiency of AAV1/6 *in vitro* and AAV9 *in vivo* for cardiac delivery and mechanistic knowledge of their action can help guide cardiac optogenetic efforts.

## INTRODUCTION

The use of adeno-associated viruses (AAVs) as transgene delivery vehicles in disease treatment requires comprehensive assessments of their performance and safety profiles. Advantages of AAVs include longterm expression, tissue tropism from 13 serotypes, and the ability to transduce both dividing and nondividing cells.^1–4^ Recent clinical trials have explored the use of AAVs in the treatment of heart failure, specifically in the upregulation of SERCA2a, a Ca^2+^ ATPase, known to be downregulated during the progression of the disease (CUPID).^5^ Although a recent CUPID phase IIb trial concluded that the delivery of SERCA2a by AAV serotype 1 did not improve symptoms of heart failure in patients, no safety issues or adverse effects were observed.^6^ As of June 2017, there have been 183 clinical trials in humans using AAV (www.abedia.com/wiley).

Concurrent to the exploration of AAV use in clinical trials, optogenetics has been rapidly developing as a promising tool in cardiac research (reviewed in ^7–10^). Optogenetics relies on the genetic modification of cells and tissues with light-sensitive opsins for precise bi-directional control of activity. The technique allows for functional manipulation of target cells/tissues with high specificity through genetic modification, in addition to the superior spatiotemporal resolution afforded by optical means.^8^ Consequently, the field of optogenetics requires highly efficient transgene delivery vehicles for cardiac applications. Such virally-mediated optogenetic manipulations are scalable by permitting the parallel investigation of many cells *in vitro* for high-throughput all-optical cardiac electrophysiology^11, 12^ or allowing cardiac applications *in vivo* across different animal species, beyond the usual mouse transgenic models.

In this study, we investigated the efficiency and mechanisms of infection of three select AAV serotypes (1, 6, and 9) with known affinity for cardiac tissue in pursuit of scalable *in vitro* and *in vivo* models for cardiac optogenetics. Our study was motivated by the inconsistency of available data and study design evaluating serotype specificity in various animal models (see Table S1 for a brief literature review). For instance, AAV9 has been shown to have highly efficient transgene delivery to the heart in the mouse and rat in a variety of studies,^13–16^ however, AAV1 and AAV6 are identified as superior for the heart in other studies.^2, 16–20^ In addition, developmental serotype specificity (i.e. preferential transgene expression in neonates versus adults) has also been suggested in studies involving dogs^21^ and rhesus macaques.^14^ A more recent study identified AAV6 as an efficient serotype for the infection of stem-cell derived cardiomyocytes.^22^ Clinically and *in vivo,* AAV-mediated gene delivery is the approach of choice, including for expression of optogenetic tools. While a number of suitable options exist for gene delivery *in vitro* other than AAV-mediated gene transfer, there is often convenience in being able to utilize the same vectors for both studies *in vitro* and *in vivo.*

We used several experimental platforms relevant to the development of viral models for cardiac optogenetics. *In vitro* we assessed serotype performance in commonly used multicellular models of cardiac tissue - neonatal rat ventricular cardiomyocytes (NRVM) and human induced pluripotent stem cell-derived cardiomyocytes (hiPSC-CM).^11^ Adult rats were also systemically infected with AAVs as their larger size compared to mice allows for *in vivo* manipulations for cardiac research, including the insertion and implantation of fiber-based devices for long-term cardiac recording and stimulation.^23^ Our final experimental platform was adult human ventricular tissue slices which may be useful as physiologically-relevant model systems for electrophysiology studies.^24^

## METHODS

Procedures involving animals were performed in accordance with institutional guidelines at both Stony Brook University and George Washington University. Procedures involving human hearts were approved by the George Washington University Institutional Review Board and informed donor consents were obtained for all tissues used in this study.

Further details of the Methods are provided in the Supplementary Material.

## RESULTS

### AAV6 Outperforms AAV1 and AAV9 *In Vitro*

Recent applications of cardiac optogenetics *in vitro,* illustrating increased-throughput electrophysiology, require the use of viral vectors to deliver genetically-encoded optical sensors or actuators.^11, 12, 30, 31^ While typically adenoviral (AdV) or lentiviral (LV) delivery has been employed in such applications, there is also interest in assessing the potential utilization of AAV vectors developed for *in vivo* applications as very few studies have been conducted in this area.^22^ To systematically quantify serotype-specific and dose-dependent AAV infection, NRVMs and hiPSC-CMs were infected with AAVs 1, 6, and 9 containing the transgene for eGFP (Figure 1). Transgene expression was cardiac specific, as eGFP was consistently co-localized in the same cells with positive immunostaining for α-actinin (Figure 1A). Although we employed strategies in the isolation of the NRVMs to reduce the presence of fibroblasts, a small number of fibroblasts are co-cultured with the cardiomyocytes, as can be seen in the non-eGFP/α-actinin-positive areas of Figure 1A. The hiPSC-CMs are, however, a purified population of cardiomyocytes and we have not observed any fibroblasts during culture.

**Figure 1.**
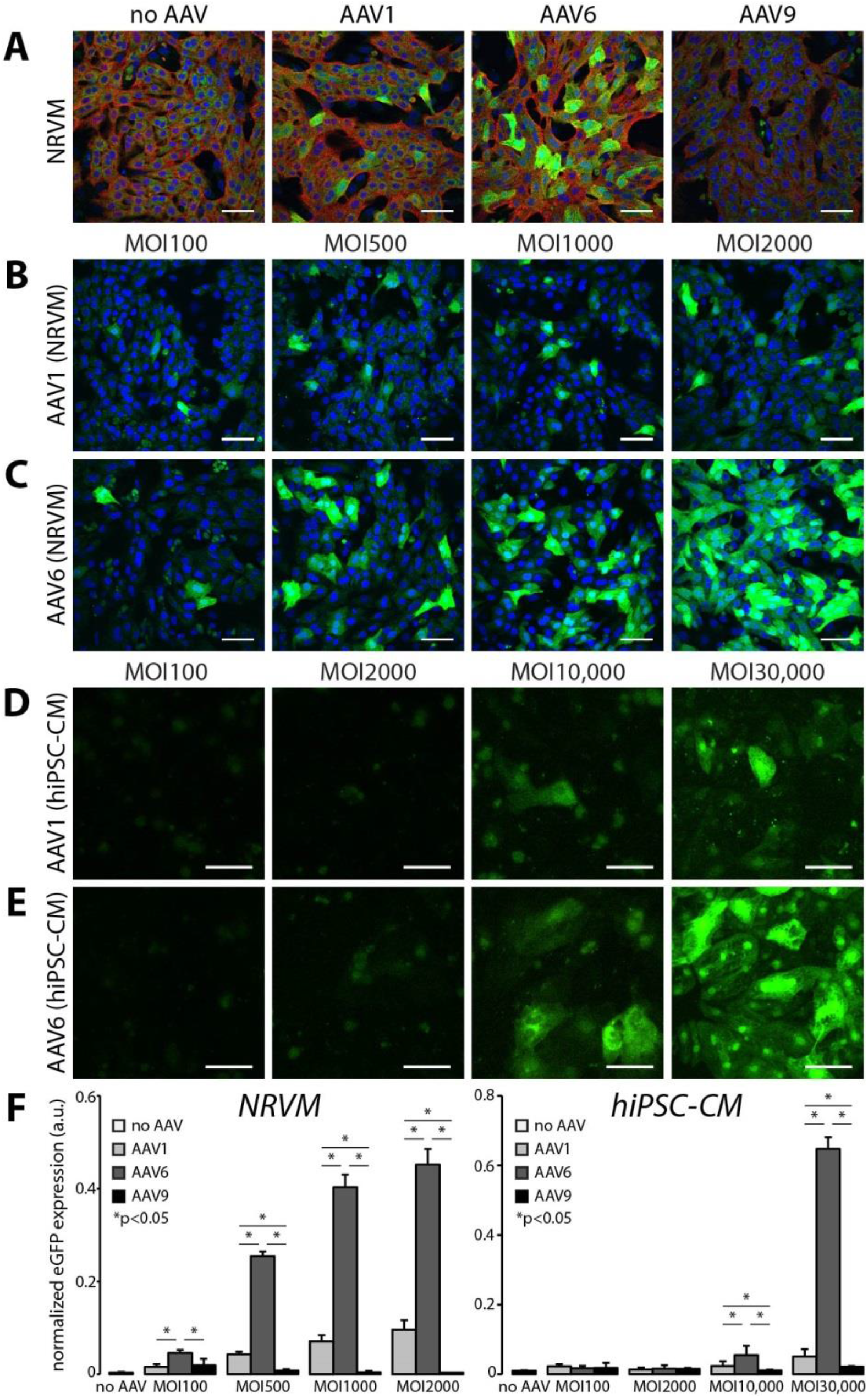
*In vitro* AAV6-mediated transgene expression is superior to the use of AAV1 and AAV9 in rat and human cardiomyocytes. **(A)** Cardiomyocyte-specific eGFP expression in NRVMs and hiPSC-CMs using AAV1, 6, and 9. AAV9-mediated expression did not exhibit levels of fluorescence above that of autofluorescence in non-infected control cells. Cell nuclei were labeled with DAPI (blue, NRVMS only), AAV-infected cells expressed eGFP (green), and cardiomyocytes were labeled with α-actinin (red). **(B,D)** AAV1-mediated and **(C,E)** AAV6-mediated eGFP expression at four viral doses five days post-infection. hiPSC-CMs required viral doses two orders of magnitude greater than NRVMs (MOI 10,000 versus MOI 100) to show threshold eGFP expression. All scale bars are 50μm and color-enhanced images are shown. **(F)** Quantification of the dose-dependent increase in eGFP expression in NRVMs and hiPSC-CMs. AAV6-mediated eGFP expression was significantly higher than AAV1-mediated expression at all viral doses. Data are presented as mean±S.E.M. (n=3-7 per group).

AAV1- and AAV6-mediated eGFP expression was dose (MOI)- dependent in both NRVMs and hiPSC-CMs (Figure 1B-E). Quantification of the AAV-mediated dose-dependency of expression showed that infection by AAV6 resulted in significantly higher transgene expression at all MOIs for NRVMs and MOIs greater than 10,000 for hiPSC-CMs. Transgene expression due to AAV1 infection was also observed, but at significantly lower levels than AAV6-mediated expression. AAV9-mediated eGFP expression was not detected at these viral doses in either cell type (Figure 1F). It should also be noted that hiPSC-CMs require viral doses two orders of magnitude greater than NRVMs (MOI 10,000 versus 100) to show baseline eGFP expression.

Viral doses greater than those shown in Figure 1 resulted in significant cell death within the monolayers. Figure S1 shows representative images of propidium iodide uptake (as a marker of dead cells) in NRVMs as a function of MOI. We quantified no significant differences in cell death across MOIs and serotypes, however, there is a trend towards increasing cell death with AAV1 infection at MOI 2000 (Figure S1D).

### AAV9 Outperforms AAV1 *In Vivo*

A prior report on cardiac optogenetics, involving systemic delivery of AAV9 encoding for the ChR2 transgene showed robust and long-lasting expression and functionality in mice.^32^ However, except for a recent brief report,^33^ to date this minimally-invasive transduction approach has not been extended to larger animals, which may be more suitable for the study of cardiac arrhythmias due to size and ease of endoscopic access.^23^ Here, systemic delivery of viral particles in the adult rat was employed through the lateral tail vein to assess the *in vivo* specificity of AAVs 1 and 9 (Figure 2). Four weeks after viral injection, excised hearts, brains, livers, and kidneys were assessed macroscopically for mCherry fluorescence (Figure 2A). AAV9-mediated infection resulted in global ventricular mCherry expression, while AAV1-mediated infection resulted in no cardiac transgene expression (with fluorescence comparable to sham viral injections) at a dose of 0.5x10^12^ viral particles per rat. Other excised organs (brain, liver, and kidney) showed little to no signs of AAV-mediated infection in all animals.

**Figure 2.**
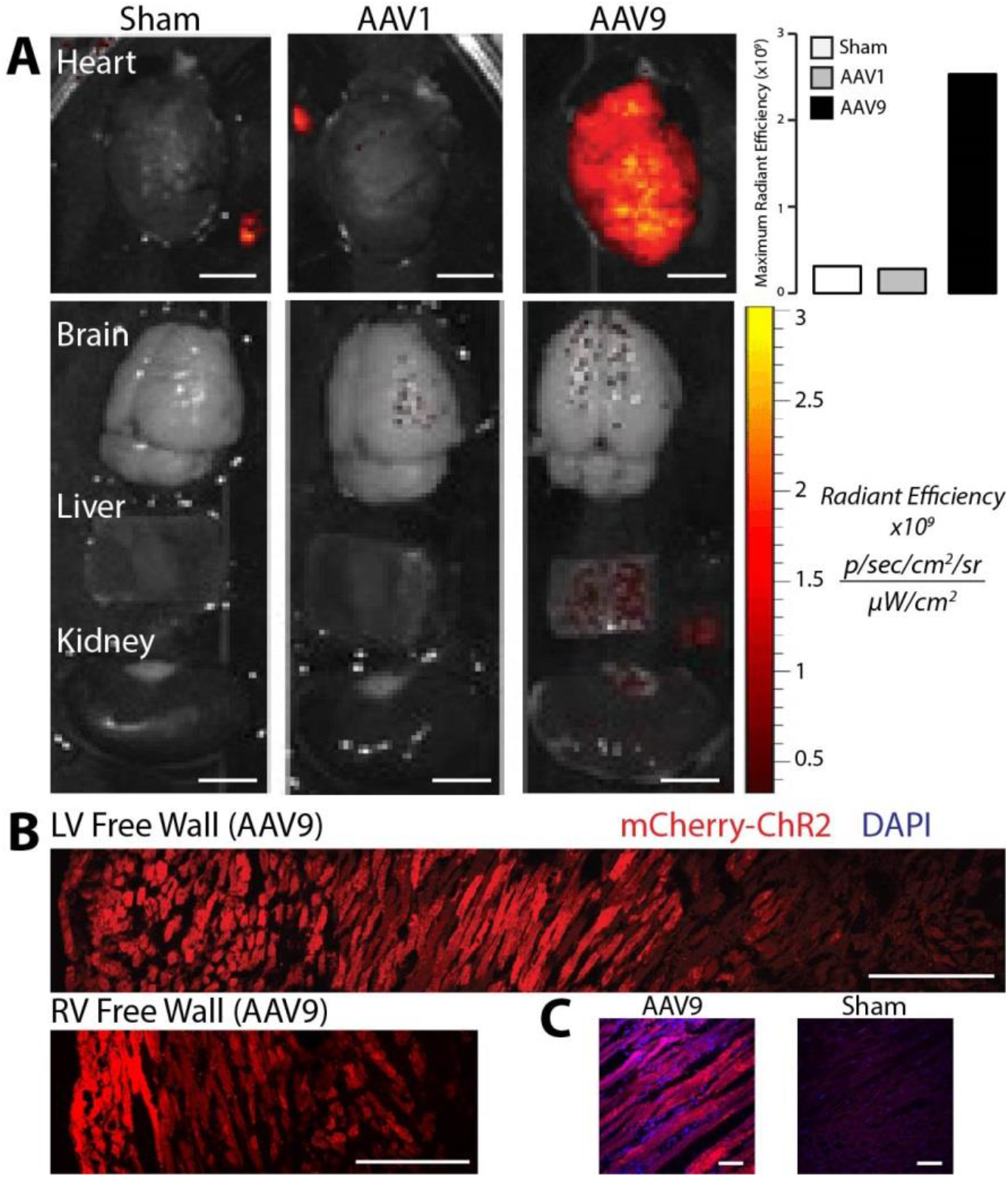
*In vivo* AAV9-mediated mCherry expression in the adult rat heart provides robust, predominantly cardiomyocyte-specific transgene delivery. **(A)** Systemic delivery of 0.5x10^12^ viral particles resulted in robust cardiac-specific expression of mCherry in four weeks using AAV9, but not AAV1 as measured using radiant efficiency. Other major organs (including brain, liver, and kidney) showed little to no signs of AAV-mediated infection. Scale bars are 500μm. **(B)** AAV9-mediated transgene delivery resulted in transmural ChR2-mCherry expression in both the LV and RV free walls. Scale bars are 250μm. **(C)** High resolution images show cardiac-specific ChR2-mCherry expression using AAV9-mediated delivery. Scale bars are 50μm.

In rats infected with AAV9, cardiac mCherry expression was robust and, not only expressed from apex to base as was observed with the macroscopic fluorescent imaging, but also expressed from the epicardium to the endocardium in both the left ventricular and right ventricular free walls (Figure 2B). Higher resolution microscopic imaging of AAV9-infected and sham hearts confirmed that observed fluorescence was not due to tissue auto-fluorescence (Figure 2C).

### AAV Serotype Infection is Mediated by Different Receptors on the Cardiomyocyte Surface

Previous studies have shown that infection by different AAV serotypes is mediated by a variety of cell surface receptors (for review see^34^). Specifically, cell surface N-linked sialic acid has been proposed as the primary receptor for AAV1 and AAV6 to infect and transduce cells.^28^ There are at least two mechanisms of AAV9-mediated cell infection/transduction involving two different receptors: terminal galactose on cell surface glycoproteins^35^ (that can be made available for AAV9 entry upon desialylation) and the 37/67 kDa laminin receptor (LamR).^36^ Figure 3 provides a visual overview of the mechanisms of infectivity of cardiomyocytes we investigated in this study.

**Figure 3.**
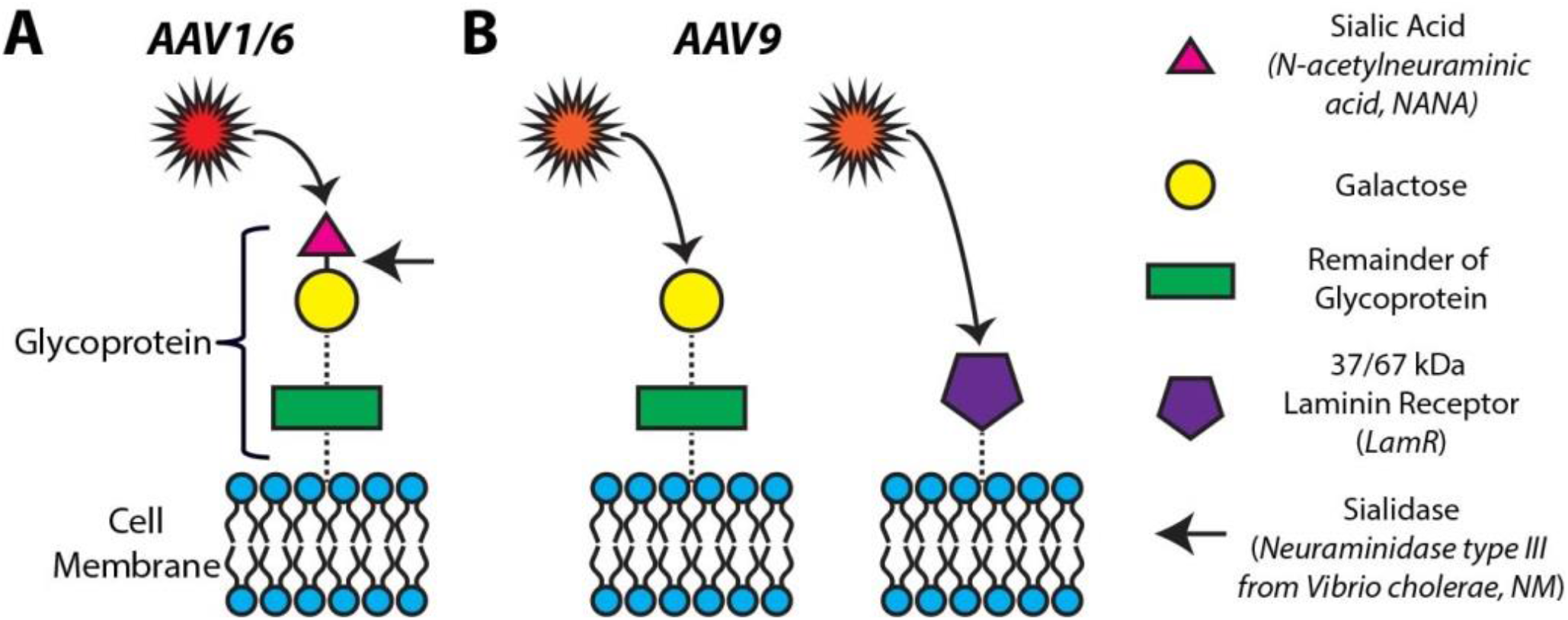
Proposed mechanisms of infectivity for AAV1, 6, and 9. **(A)** Cell surface N-linked sialic acid has been proposed as the primary receptor for AAV1 and 6 to infect and transduce cells. The removal of sialic acid by neuraminidase (targeting the portion of the glycoprotein indicated by the arrow) is expected to block the AAV1,6-mediated transduction of cells. **(B)** AAV9-mediated cell infection/transduction has been attributed to two receptors: terminal galactose on cell surface glycoproteins (left panel) and the 37/67 kDa laminin receptor (right panel).

In order to probe the mechanisms of our differential observations of AAV serotype specificity *in vitro* (Figure 1) and *in vivo* (Figure 2), we explored the roles of both sialic acid and LamR in AAV-mediated transgene expression. As indicated in Figure 3A by the arrow, we hypothesized that the removal of sialic acid by neuraminidase (NM) would block AAV1- and AAV6-mediated infection of cells. On the other hand, the same removal of sialic acid would also free up terminal galactose on the cell surface thus enhancing AAV9-mediated infection (Figure 3B, left panel). Similarly, AAV9 infection would be enhanced by the presence of LamR on the cell surface (Figure 3B, right panel).

### *In Vitro* Desialylation Modulates AAV-Mediated Gene Expression

Treatment of both NRVMs and hiPSC-CMs with NM, a broad spectrum sialidase, to remove cell surface sialic acid significantly reduced eGFP expression via AAV1 and AAV6 (Figure 4). In NRVMs, AAV1-mediated eGFP expression was completely abolished by 25mU/mL NM, whereas AAV6-mediated expression was reduced to the point where only a few individual cells were eGFP-positive (Figure 4A,B). AAV9-mediated eGFP expression was unaffected in NRVMs as we did not observe the purported enhanced entry of AAV9 (Figure 3B) even at higher NM doses.

**Figure 4.**
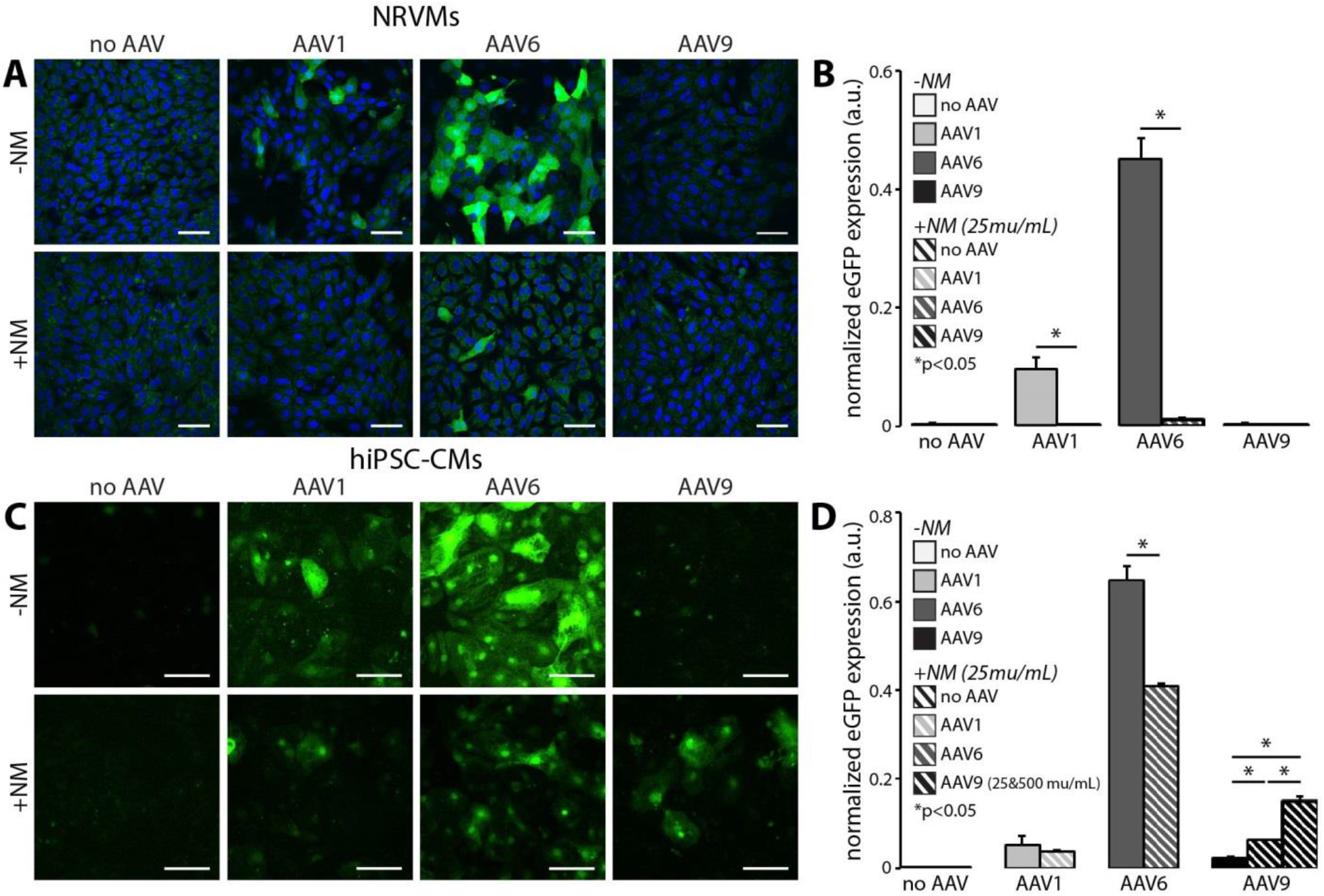
*In vitro* desialylation modulates AAV-mediated eGFP expression in NRVMs and hiPSC-CMs. Cardiomyocyte-specific eGFP expression with (+NM, 25 mU/mL) and without (-NM) neuraminidase treatment prior to viral infection in **(A)** NRVMs at MOI 2000 and **(C)** hiPSC-CMs at MOI 30,000. Cell nuclei were labeled with DAPI (blue, NRVMs only) and AAV-infected cells expressed eGFP (green). All scale bars are 50μm and color-enhanced images are shown. Quantification of eGFP expression with and without desialylation in all three serotypes in **(B)** NRVMs and **(D)** hiPSC-CMs. eGFP expression mediated by AAV1 and 6 significantly decreased in both cell types, whereas transgene expression mediated by AAV9 significantly increased in hiPS-CMs only. Application of a higher dose of NM (500 mU/mL) in hiPSC-CMs infected with AAV9 resulted in even greater eGFP expression. Data are presented as mean±S.E.M. (n=3-7 per group).

The same dose of NM in hiPSC-CMs never completely eliminated transgene expression, but AAV1 and AAV6-mediated infection was significantly reduced (Figure 4C,D), similar to the effect observed in NRVMs and in line with the predictions from Figure 3A. Interestingly, the application of NM to hiPSC-CMs in combination with AAV9 infection significantly increased transgene expression (Figure 4C,D). The additional application of 20x our standard NM dose (500 mU/mL), resulted in a further increase in AAV9-mediated gene expression, beyond that of AAV1-mediated expression without NM (Figure 4D), presumably by exposing terminal cell surface galactose for infection by AAV9, as illustrated in Figure 3B, and in contrast to our findings in NRVM.

### *In Vivo* 37/67 kDa Laminin Receptor (LamR) Expression in the Adult and Neonatal Rat Heart and Human Ventricular Tissue Slices

The observed discrepancies in AAV serotype-mediated transgene expression *in vitro* (where AAV6 was most efficient) and *in vivo* (where AAV9 was most efficient) were further elucidated by investigating the presence of LamR which is purported to be a cell surface receptor for AAV9, as previously discussed. Our data show that LamR is not present *in vitro* in monolayers of NRVMs, nor in hiPSC-CMs (Figures 5A, S3A), but appears to be globally present *in vivo* in both the adult and neonatal rat heart (Figures 5B, S2B). Wild-type HeLa cells served as our *in vitro* positive control (Figure 5A) and tissue sections of breast carcinoma served as our *in vivo* positive control (Figures 5B, S2A). Additional staining without the primary antibody showed that *in vivo* the secondary antibody did not yield any non-specific staining (Figure 5C).

**Figure 5.**
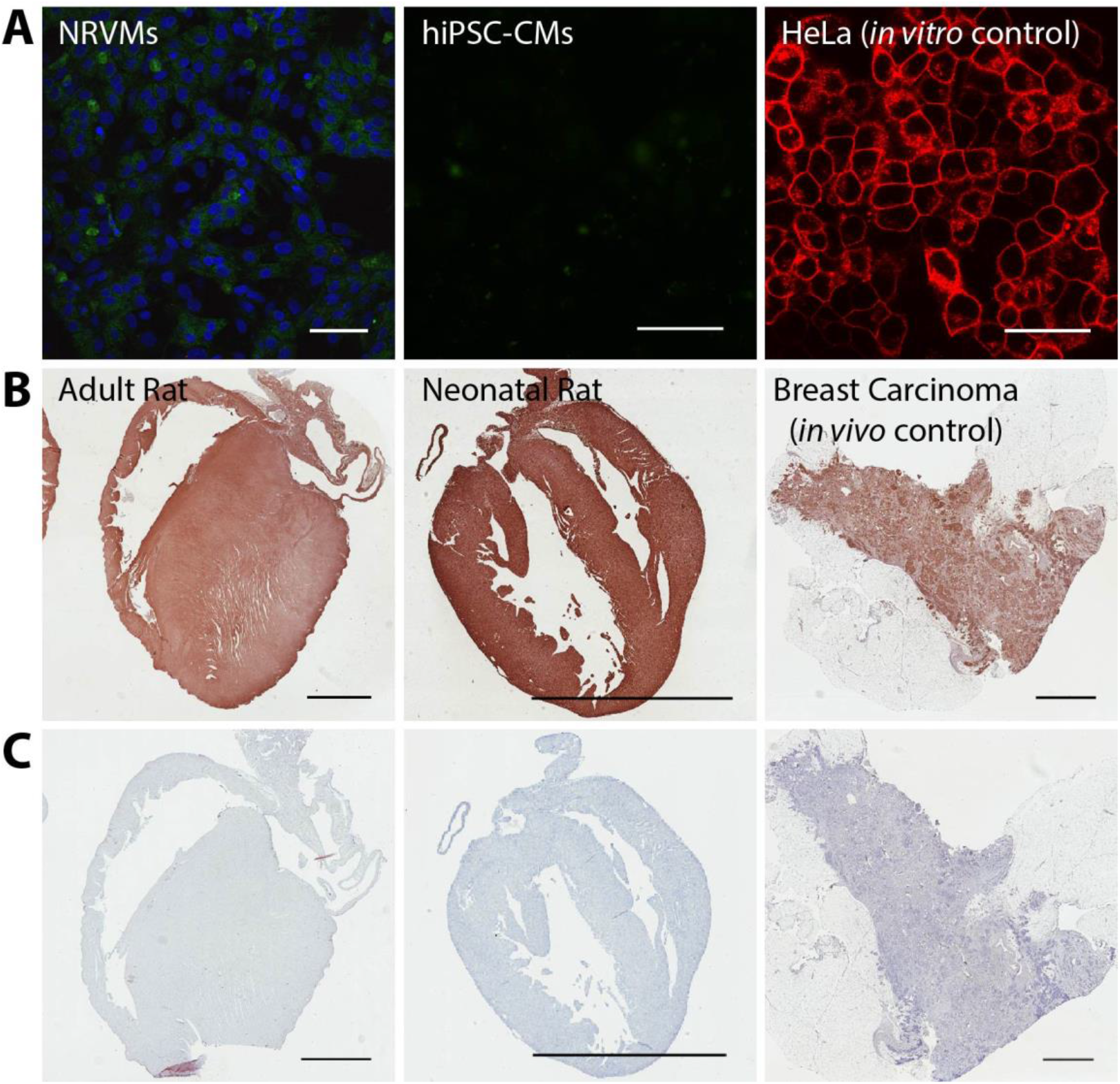
The 37/67 kDa laminin receptor (LamR) is expressed in the rat heart, but not in cultured NRVMs and hiPSC-CMs. **(A)** Negative *in vitro* immunostains of NRVMs and hiPSC-CMs for LamR. Concurrent immunostaining of HeLa cells, serving as a positive *in vitro* control for LamR. Scale bars are 50μm. **(B)** Positive immunostains of adult and neonatal rat hearts. Concurrent immunostaining of breast carcinoma tissue, serving as a positive *in vivo* control for LamR. **(C)** Negative controls of tissue (stained with no primary LamR antibody) showed no contribution to the positive stain by nonspecific secondary antibody staining. Scale bars in B and C are 3mm.

Extending these observations to the adult human heart, we looked at LamR expression in human ventricular tissue slices post-extraction (Figure 6A), as well as after 36 hours in culture (Figure 6B). The fresher heart slices showed very minimal LamR expression, in contrast to our results with the fresh rodent hearts. This could be due to long cross-clamp to fixation time (5 to 12 hours for the two hearts used here). After 36 hours in culture, unlike our observations in cultured cells, there was some recovery of LamR expression in the human heart tissue slices, which remained lower than that seen in the rodent hearts. Treatment of the tissues with NM (500mU/mL) during culture did not alter LamR expression (Figure S2C).

**Figure 6.**
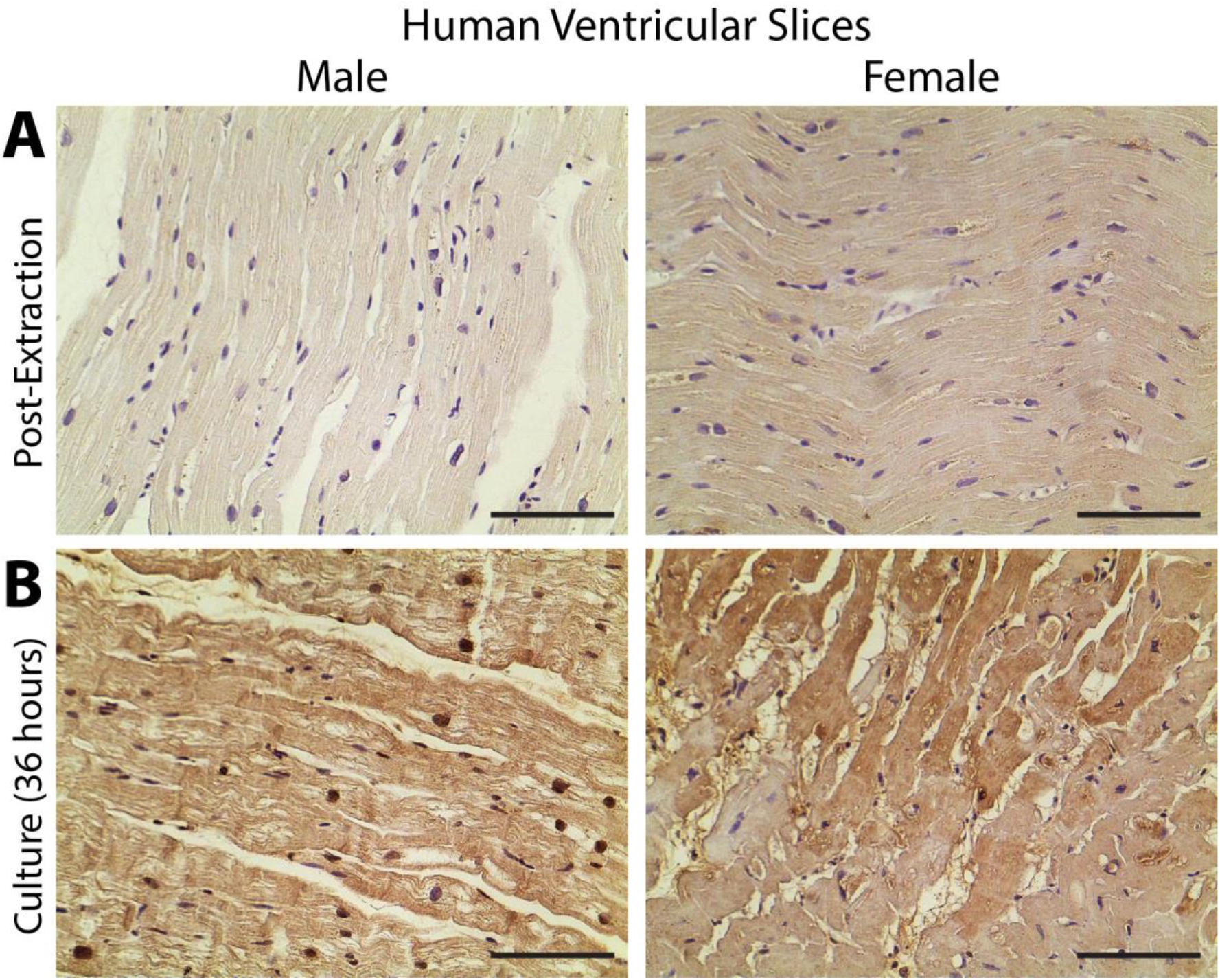
The 37/67 kDa laminin receptor (LamR) is expressed in human ventricular tissue slices. Positive immunostains of male (left) and female (right) ventricular slices **(A)** post-extraction with minimal expression and **(B)** after recovery of 36 hours in culture. The time between cross-clamp of the heart and fixation in **(A)** was >12 hours in the male specimen and ~5 hours in the female specimen. This difference is reflected as lower LamR expression in the male heart slice immediately post-extraction **(A)** which then recovered to expression levels similar to the female slice 36 hours later **(B)**. Scale bars are 100μm.

Given our LamR expression data in NRVMs, hiPSC-CMs, rat hearts, and human tissue slices, it is important to note that the expression of LamR is dynamic and significantly affected by the tissue/culture environment. Specifically, we have observed the paucity of LamR expression in the *in vitro* environment with isolated cells (Figures 5A, S3) compared to its robust presence *in vivo* (Figures 5B, S2A) and *in situ* in intact tissues (Figures 6, S2C).

### TGF-β1 Treatment Does Not Significantly Affect AAV9-Mediated Gene Expression

Since the presence of LamR was not detected *in vitro* in NRVMs and hiPSC-CMs, we followed up on an earlier report^29^ and pre-treated the monolayers with TGF-β1 (10ng/mL for 24 hours) in an attempt to increase LamR expression and facilitate AAV infectivity. Our data, however, shows that TGF-β1 application does not significantly increase the expression of LamR (Figure S3A,B) and minimally increases AAV9-mediated eGFP expression (Figure S4).

## DISCUSSION

We investigated *in vitro* and *in vivo* AAV serotype specificity in rat and human models suitable as scalable experimental platforms for cardiac optogenetics. Different optimal serotypes were identified for *in vivo* and *in vitro* use. Namely, *in vitro* AAV6-mediated transgene expression was superior to AAV1,9-mediated delivery due to the presence of cell surface N-linked sialic acid (Figures 1 and 4). The subsequent enzymatic removal of sialic acid significantly reduced or abolished AAV6- and AAV1-mediated gene delivery, independent of cell type. Interestingly, the same desialylation enhanced AAV9-mediated expression but only hiPSC-CMs (Figure 4). In contrast, *in vivo* serotype specificity in the adult rat favored delivery by AAV9, likely mediated by the presence of cell surface LamR (Figures 2 and 5), which appears ubiquitous in the intact heart and to a certain degree in recovered explanted human cardiac tissue, but could not be found in cultured cardiomyocytes.

### Viral Delivery of Optogenetic Tools *In Vitro* for High-Throughput Cardiac Electrophysiology

The growing use of optogenetics in cardiac applications motivated our search for optimized parameters for the optical control of the heart under various experimental conditions. One such application is the development of high-throughput all-optical electrophysiology for drug screening and cardiotoxicity testing.^11, 12^ The current study revealed that the environment (i.e. *in vitro* versus *in vivo)* is of great importance with regard to preferential serotype specificity. Cultured cardiomyocytes tend to lose cell surface receptors (LamR) critical to mediating *in vivo* AAV9 infection (Figure 5), although those receptors are present *in situ* in cultured explanted human cardiac tissue. Consequently, the *in vitro* environment is very different than that of the intact heart, requiring different means for efficiently inscribing optical control.

AAV6-mediated transgene delivery resulted in acceptable expression levels, similar to those of our previous studies using adenoviral delivery.^11, 26, 27^ However, the required dose (MOI) was orders of magnitude higher (Figure S5), and the time required for transgene expression with AAV is not optimal for primary cells. Here AAV infection required five days for the cells to reach peak transgene expression (data not shown), whereas in our previous studies >95% of NRVMs and hiPSC-CMs expressed ChR2 within 24-48h using an adenovirus (Figure 7A).^11, 26^ *In vitro* optical control was confirmed using all-optical electrophysiology (combining optical mapping by voltage-and calcium-sensitive dyes with simultaneous optogenetic stimulation).^11^

**Figure 7.**
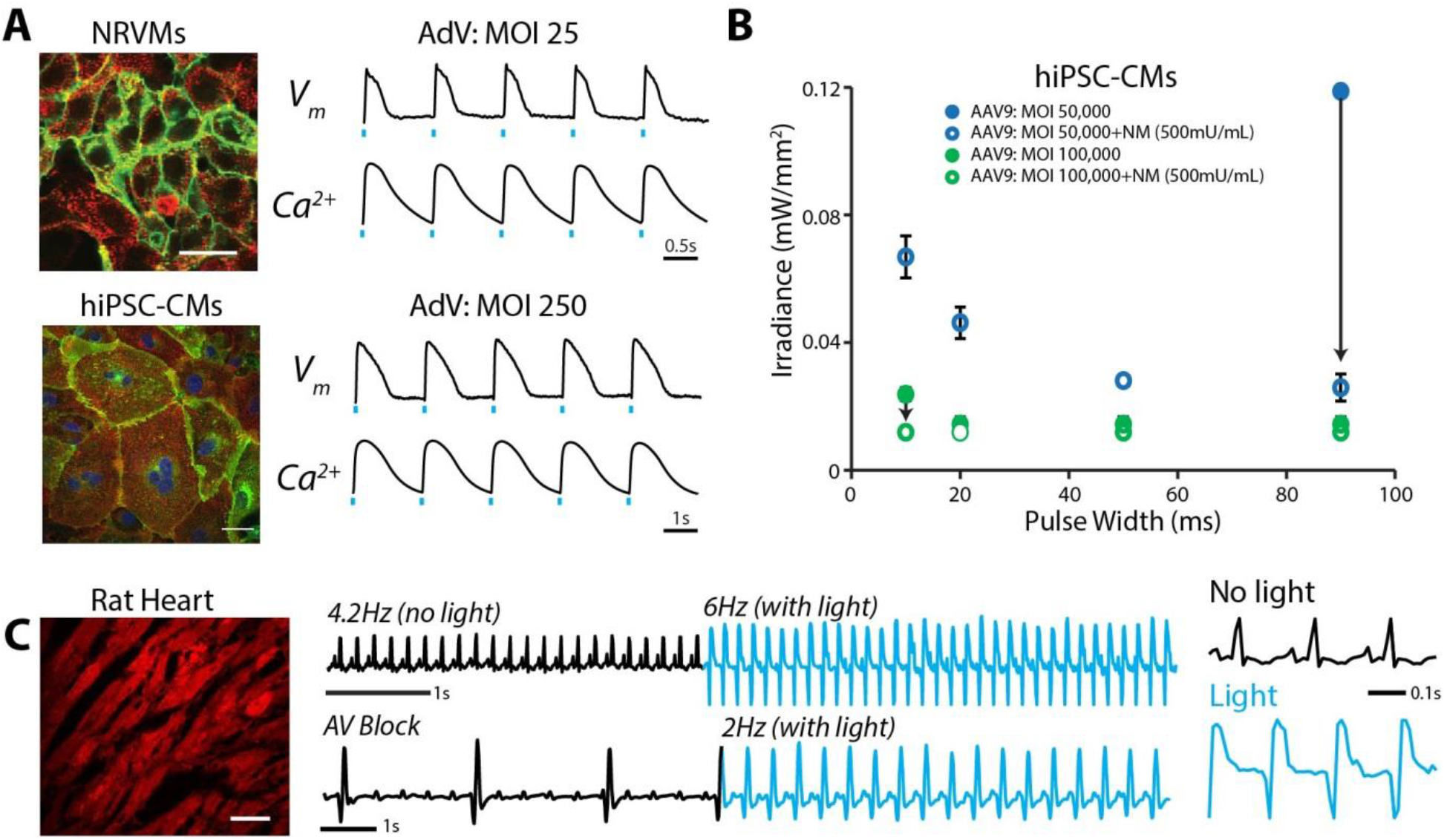
Robust *in vitro* and *in vivo* optogenetic control of the heart. **(A)** Adenoviral (AdV)-mediated ChR2-eYFP expression and functional measurements in NRVMs (MOI 25) and hiPSC-CMs (MOI 250) two days post infection. Functional measurements were acquired using voltage-(di-4-ANBDQBS) and calcium-(Rhod4) sensitive dyes and example traces with optical pacing are shown. Cell nuclei were labeled with DAPI (blue) and AdV-infected cells expressed eYFP (green). Alpha-actinin staining (red) showed the cardiospecificity of the ChR2-eYFP infection. **(B)** Strength-duration curves for AAV9-mediated ChR2 expression in hiPSC-CMs. Conditions for infection included MOIs of 50,000-100,000 and NM applications of 500 mU/mL. Black arrows show the effect of NM treatment on lowering irradiance (mW/cm^2^) requirements. Data are presented as mean±S.E.M. (n=3 per group). **(C)** AAV9-mediated ChR2-mCherry expression in the intact adult rat heart after 4 weeks results in optically-sensitive myocardium *in situ* (left panel). A 0.8mm diameter optical fiber was used to optically control electrical activity from the LV free wall as recorded using ECG (middle panel). Optical pacing resulted in an increased heart rate, as well as significant morphological changes in the QRS complex (right panel). Spatial scale bars are 50μm in **B** and **C**. Temporal scale bars are as indicated.

Although *in vitro* AAV9-mediated transgene delivery was deemed less optimal than AAV1,6-mediated delivery (Figure 1), successful expression of ChR2-mCherry using AAV9 was achieved under very specific conditions, as hypothesized and explored in this study (Figure 7B). A pre-treatment of hiPSC-CM monolayers with 500mU/mL NM (20x the dose required to cause desialylation, Figure 4), followed by AAV9 infection at very high MOIs of 50,000-100,000 (5-10x the minimum dose for baseline transgene expression, Figure 1) resulted in optogenetic responsiveness. In all four cases (MOI 50,000±NM and MOI 100,000±NM), ChR2-mCherry was expressed resulting in an optically-sensitive cardiac syncytium. However, at the lower concentration of MOI 50,000 only (no NM) and relevant low-light stimulation, only 1 out of 3 samples was optically excitable. As illustrated in Figure 7B, the strength-duration relationship showed the effect of NM treatment on improving optical responsiveness as compared to infection alone (black arrows). Despite successful transgene expression, infection at such high MOIs resulted in significant cell death (data not shown).

The emergence of human stem-cell derived cardiomyocytes and their combination with genetically-encoded sensors and actuators^11, 12, 30, 31^ has prompted a closer look at the performance of various viral vectors, including AAVs^22^ due to the convenience of sharing the usage of such vectors for both *in vivo* and *in vitro* applications. The results presented here, showing preferential infectivity of cardiomyocytes *in vitro* (AAV6>>AAV1>>AAV9), are consistent with a recent report in human stem-cell derived cardiomyocytes. Interestingly, we find that the ease of viral infection in the *in vitro* environment seems to be dependent on two major factors: the viral vector itself (AAV, adenovirus, or lentivirus) and the state of differentiation of the target cell (Figure S5). In our experience, primary cardiomyocytes are the easiest to infect (i.e. requiring the lowest viral doses for >80% cell transgene expression) using AdV^26, 27^ and AAV (explored in this study). iPSC-CMs require 10-100x increased viral doses compared to primary cardiomyocytes for the same efficiency of expression, and the presumably least differentiated cells, cardiac fibroblasts, require the highest viral doses,^37^ although we have not tested AAVs on the latter cell type (Figure S5). Similar observations have been reported for pluripotent stem cells before and after differentiation into cardiomyocytes.^22^ While AAV delivery appears sub-optimal for *in vitro* use (compared to LV or AdV application), our dissection of the mechanism of viral entry suggests some strategies to improve infectivity with select AAVs, e.g. desialylation enhances AAV9-mediated entry, while sialic acid on the cardiomyocyte surface promotes AAV6 entry.

### Evolution of AAV as an *In Vivo* Gene Therapy Tool

Optogenetics in the intact organism requires the genetic modification of cells and tissues, and hence it necessitates the development of efficient, safe tools for gene therapy. Methods for non-viral transfer of genetic material, including electroporation, ballistic DNA transfer, and cationic lipid-based gene transfer, are known to be less efficient and the persistence of transgene expression is short-lived.^38^ Therefore, viral transfer of genetic material through the use of adenoviruses, lentiviruses, and AAVs is desirable. AAVs are preferred due to their comparatively low immunogenicity.^39^ *In vivo,* AAV-mediated transgene delivery has been used for cardiac optogenetics in rodent hearts.^32, 33, 40^ AAV9-mediated expression of ChR2 in the mouse heart yielded highly efficient and cardiac-specific transduction when applied by a minimally-invasive systemic route;^32^ a recent report used a similar delivery but with a very high dose of cardiac-specific viral vector in the rat.^33^ Direct cardiac injections of AAV9 encoding for the ChR2 transgene also resulted in optical responsiveness of the rat heart.^40^ However, systemic delivery is preferred not only because of its minimally-invasive nature (and hence, suitability for translation), but also because of better uniformity of expression.^32, 41^ Here, we show systemic delivery and successful expression of ChR2-mCherry in the adult rat heart four weeks after viral injection using AAV9 with a generic promoter (Figure 7C). Optical sensitivity was confirmed by rate and QRS morphology changes in the ECG, when using an optical fiber to deliver light to the left ventricle in the open chest of the anesthetized rat.

AAV serotypes 1, 6, and 9 have shown different degrees of gene transfer to the heart (Table S1). In addition to the AAV tissue tropism, the use of cardiac-specific promoters, such as cardiac troponin T (cTnT), has been shown to further increase specificity, however the level of expression derived from tissue-restricted promoters may not be as high as from ubiquitous viral promoters.^41^

Of critical importance in the use of AAV serotypes for optimized gene therapy applications is consideration of the mechanism of infectivity. In this study, we not only identified optimal serotypes for *in vitro* and *in vivo* use, but also explored the mechanism of infection and fundamental differences between experimental platforms. Specifically, our results are consistent with AAV9 infection being mediated by either terminal galactose (Figure 4, only in hiPS-CMs) or LamR (Figure 5). Identification of unique cell surface receptors in the heart and other organs will continue to drive the design of truly optimized AAV serotypes for cardiac application, such as optogenetics, and beyond.

## ACKNOWLEDGMENTS

We acknowledge the technical support provided by the Research Pathology Core Laboratories at both Stony Brook University and George Washington University, as well as the assistance of veterinary personnel in the Division of Laboratory Animal Resources at Stony Brook University and the Animal Research Facility at George Washington University. We would also like to acknowledge Dr. Igor Efimov and Jaclyn Brennan in the procurement and culture of the human tissue slices.

